# A novel approach reveals that HLA class 1 single antigen bead-signatures provide a means of high-accuracy pre-transplant risk assessment of acute cellular rejection

**DOI:** 10.1101/433318

**Authors:** Nicole Wittenbrink, Sabrina Herrmann, Arturo Blazquez-Navarro, Alessandro Gulberti, Chris Bauer, Eric Lindberg, Petra Reinke, Birgit Sawitzki, Oliver Thomusch, Christian Hugo, Nina Babel, Harald Seitz, Michal Or-Guil

**Affiliations:** Systems Immunology Lab, Department of Biology, Humboldt University Berlin, Germany; Fraunhofer Institute for Cell Therapy and Immunology, Bioanalytics und Bioprocesses, Potsdam, Germany; Berlin-Brandenburg Center for Regenerative Therapies (BCRT), Berlin, Germany; MicroDiscovery GmbH, Berlin, Germany; Department of Nephrology and Internal Intensive Care, Charité University Medicine Berlin, Campus Virchow Clinic, Berlin, Germany; Molecular Immune Modulation, Institute for Medical Immunology, Charité University Medicine Berlin, Campus Virchow Clinic, Berlin, Germany; Klinik für Allgemein- und Viszeralchirurgie, Universitätsklinikum Freiburg, Freiburg, Germany; University Hospital Carl Gustav Carus, Dresden University of Technology, Dresden, Germany; Medical Clinic I, Marien Hospital Herne, Ruhr University Bochum, Herne, Germany.

**Keywords:** renal transplantation, acute cellular rejection, pre-transplantation risk assessment, anti-HLA-1 antibodies, single HLA antigen bead assay, immune signatures, machine learning

## Abstract

Acute cellular rejection (ACR) is associated with complications after kidney transplantation, such as graft dysfunction and graft loss. Early risk assessment is therefore critical for the improvement of transplantation outcomes. In this work, we retrospectively analyzed a pre-transplant HLA antigen bead assay data set that was acquired by the e:KID consortium as part of a systems medicine approach. The data set included single antigen bead (SAB) reactivity profiles of 52 low-risk graft recipients (negative complement dependent cytotoxicity crossmatch, PRA<30%) who showed detectable pre-transplant anti-HLA 1 antibodies. To assess whether the reactivity profiles provide a means for ACR risk assessment, we established a novel approach which differs from standard approaches in two aspects: the use of quantitative continuous data and the use of a multiparameter classification method. Remarkably, it achieved significant prediction of the 38 graft recipients who experienced ACR with a balanced accuracy of 82.7% (sensitivity=76.5%, specificity= 88.9%). The resultant classifier achieved one of the highest prediction accuracies in the literature for pre-transplant risk assessment of ACR. Importantly, it can facilitate risk assessment in non-sensitized patients who lack donor-specific antibodies. As the classifier is based on continuous data and includes weak signals, our results emphasize that not only strong but also weak binding interactions of antibodies and HLA 1 antigens contain predictive information.

## 1. Introduction

The efficacy of immunosuppressive therapy in kidney transplantation has steadily increased over the last decade. As a consequence, the incidence of acute rejection (AR) episodes has decreased and short-term graft survival rates have improved (1,2). However, long-term transplant outcomes are still poor and episodes of AR are known to significantly exacerbate long-term outcomes (2,3). AR is associated with long-term complications, such as graft dysfunction and reduced graft survival and AR prevention continues to be a main focus in the design of new therapeutic strategies for renal transplantation (4–7). The most common form of AR is acute cellular rejection (ACR) (8). ACR is a T cell cytotoxic immune response against the graft, leading to inflammatory cell infiltration with tubulitis and, eventually, damage of the donor tissue (9,10). The positive outcome of ACR if treated early, as well as its potentially irreversible damage, render it particularly relevant for prevention research (10,11). Regarding non-invasive diagnostics, a number of studies have obtained good results using tissue, blood or urine markers (11–18). For early risk assessment, the large majority of models are donor-dependent, as they either employ measurements from the early post-transplantation period or utilize donor–derived data (e.g. from crossmatch tests) (19,20,29,30,21–28). The most common approach for pre-transplant risk assessment relies on the characterization of HLA antibodies in recipient serum samples by solid phase single HLA antigen bead (SAB) assay (22–27,31). The assay facilitates detection and identification of anti-HLA antibody specificities and provides a method for monitoring the development of donor-specific antibodies (DSA). The detection of DSA through SAB assays is a well-established method for antibody-mediated rejection (ABMR) pre-transplantation risk assessment, but not for ACR (22–28,32).

Approaches for risk assessment of ACR do not employ DSA for the prediction – as both patients with or without DSA experience episodes of ACR – but other risk markers, such as soluble CD30 levels or panel of reactive T cells (21,33–35). However, the inspection of SAB serum antibody reactivity profiles (irrespective of DSA status) may provide a means to an ACR risk assessment tool for two reasons: (1) serum antibody binding profiles against antigen/protein libraries are generally powerful in discriminating between different health or disease conditions (36–39), and (2) antibody-mediated mechanisms have been shown to be involved in the T cell-mediated initiation, perpetuation, and progression of graft injury (40,41).

In this work, as part of an exploratory study, we present a classifier achieving high-accuracy pre-transplant risk assessment of ACR. Remarkably, this classifier is based on continuous non-thresholded HLA 1 SAB data and does not rely on donor-specific HLA typing.

## 2. Experimental procedures

### 2.1. Patient population and monitoring

615 adult kidney transplant recipients were enrolled in the randomized, multicenter diagnostic trial Harmony (EudraCT-Nr. 2007-006516-31) (4). Patients were treated with a quadruple (arm A) or triple (arms B and C) immunosuppressive therapy as described before (4). The immunosuppressive therapy included induction with either monoclonal IL-2R antibody basiliximab (arms A and B) (Simulect^®^, Novartis) or rabbit ATG (arm C) (Thymoglobulin^®^, Sanofi). Maintenance immunosuppression consisted of tacrolimus (Advagraf^®^, Astellas) and mycophenolate mofetil (MMF) with (arm A) or without steroids (arms B and C) (4). All transplantations were of low immunological risk, with recipient PRA scores ≤ 30% and no detectable DSA prior to transplantation (complement-dependent cytotoxicity crossmatch) (4). Further inclusion and exclusion criteria can be found in Thomusch *et al.* 2016 (4). Suspected episodes of acute rejection were confirmed through biopsy according to the Banff criteria of 2005 (42). For the e:KID project, which aims at early risk assessment of ACR by following a systems medicine approach (43,44), 157 recipients were retrospectively monitored for the presence of HLA antibodies in blood serum on day 0 (pre-transplantation). All patients who experienced ACR (borderline or Banff class 1 or higher) in the first year were assigned to the ACR group (N=77). The control group included all patients who neither experienced a rejection episode nor other serious adverse events (N=80). The study was carried out in compliance with the Declaration of Helsinki and Good Clinical Practice. All participants provided written informed consent prior to inclusion into the study.

### 2.2. HLA antibody detection by Luminex multiplex bead assay

Screening for HLA class 1 and class 2 antibodies was performed using a MAB assay (LABScreen^®^ Mixed Kit, One Lambda, Canoga Park, CA, USA). All sera that tested positive and a random subset of negative sera were subject to SAB assays to identify antibody specificities (LABScreen Single Antigen HLA Class I kit and/or LABScreen Single Antigen HLA Class II kit, One Lambda). Both MAB and SAB were performed according to the manufacturer’s instructions. Briefly, following heat-inactivation at 56°C for 30 min and clearance from debris (0.22 μm filter), 20 μl of undiluted serum was added to 3 μl of the LABScreen bead mix and incubated for 30 min in the dark at room temperature. After a washing step in 1× LABScreen wash buffer, the bead mix was incubated with 100 μl of a 1:100 dilution of the PE-conjugated goat anti-human IgG detection antibody for 30 min in the dark at room temperature. After a final washing step in 1× LABScreen wash buffer, data acquisition was performed using a FLEXMAP3D Analyser in combination with xPONENT software version 4.1 (Luminex Corporation, Texas, USA).

### 2.3. Conventional HLA data processing and analysis

Key to the conventional method for HLA data processing is the binarization of the continuous xPONENT median fluorescence intensity (MFI) raw data by means of a MFI theshold (1=presence of reactivity, 0=absence of reactivity). For generation of binary data, raw MFI data were normalized to an in-house negative control serum (MAB) or the One Lambda negative serum OLI.NS (SAB). In case of MAB data, a bead was considered positive if its normalized background ratio exceeded 3. For binarization of SAB data, we used both a fixed threshold of 1000 MFI and an individually adjusted MFI threshold. In case of the latter, a bead was considered positive when its baseline normalized MFI exceeded 30% of the MFI of the bead showing the highest strength in reaction. Single parameter prediction performance of binarized HLA class I SAB data was assessed using Fisher’s exact test. In the case of binarized HLA class reactivity screening data (MAB), study groups were compared using Fisher’s exact test.

### 2.4. Experimental design and statistical rationale: Novel strategy for HLA data processing and analysis

In this work we applied a novel approach to the HLA data sets that does not rely on MFI-thresholding. Key to it is the rank-normalization of the continuous xPONENT median fluorescence intensity (MFI) raw data. To assess the predictive potential of the rank-normalized data, we performed multiparameter classification using an R version 2.15.0 implementation of the Potential Support Vector Machine (P-SVM) algorithm. To assess the predictive performance of the classifier, we followed a leave-one-out cross-validation approach. The latter provides a well-established solution to assess a classifiers’ predictive performance for high-dimensional, low sample size data sets as ours (45). To rate predictive performance, we used the statistical measures sensitivity, specificity and balanced accuracy (BACC). To assess the statistical significance of the predictive performance, we used random class label-permutation testing.

The significance level was set at p<0.05. The training of the P-SVM classifier involves two hyperparameters: *C* (cost) and ε (regularization). In case of classification of SAB data, we performed grid search for the hyperparameter space ε={0.25, 0.5, 0.75, 1} and *C*=*{1, 6}*; the hyperparameter space for MAB data-based classification was ε={8, 9, 10, 11} and *C*=*{1, 6}*. True classification and p-value estimation were always carried out for the same grid of hyperparameters. To further specify the performance of classificators, receiver operating characteristic (ROC) curve analysis was performed. The numerical scores (decision values) that form the basis of P-SVM class identity label assignment were extracted, sorted in increasing order and used as decision boundaries. For each boundary, both sensitivity and specificity were estimated. AUC was calculated based on Mann-Whitney U statistics (46).

### 2.5. Statistical analyses

To assess whether the two study groups (control/ACR) differed in any of the baseline population characteristics, Mann-Whitney U test (metric variables), Pearson’s chi-squared or Fisher’s exact test (categorical variables) were applied. A p-value < 0.05 was considered statistically significant. Variables are described with mean±standard deviation or median [interquartile range (IQR)].

## 3. Results

### 3.1. Characteristics of the graft recipients included in the study

Pre-transplant HLA assay data were retrospectively analyzed as part of a systems medicine approach towards early risk assessment of ACR (43,44). The investigated study group comprised all kidney transplant recipients enrolled in the Harmony trial (N=615) who experienced at least one ACR or borderline ACR event in the first year (N=77) and all transplant recipients who experienced no serious adverse events (N=80). Median time to the first ACR event was 20.5 days (range=4-373 days) (Supplemental Figure 1). Demographics and clinical characteristics of the study groups are summarized in Supplemental Table 1.

Pre-transplantation HLA-1 and HLA-2 MAB data were available for N=63 recipients of the ACR group and N=54 recipients of the control group (for demographic and clinical characteristics, see Supplemental Table 2). Additionally, HLA-1 SAB data was measured for all those patients who tested positive for HLA-1 MAB screening (21 ACR + 13 control) and a random subset of patients who tested negative (13 ACR + 5 control). In total, pre-transplantation HLA-1 SAB data were available for N=34 recipients of the ACR group and N=18 of the control group. Due to the higher sensitivity of SAB assay compared to MAB, the former assay was considered a better candidate for ACR risk assessment.

Demographic and clinical characteristics of the SAB ACR (N=34) and the SAB control group (N=18) were compared and are summarized in Table 1. The majority of patients was male, received their first kidney transplant and had a deceased donor. There were no significant differences between the study groups for the above mentioned characteristics as well as immunosuppressive therapy. However, the mean age of patients in the ACR group (54.9±11.0) was significantly higher than for the control group (51.6±11.6; p=0.04; Mann-Whitney U test). With respect to HLA mismatches, a significant difference was found for HLA-DR (p = 0.03; Pearson’s chi-squared test), with an elevated frequency of patients with two mismatches in the ACR group (32.4% vs. 11.1%). No significant differences were found for HLA-A or HLA-B. There were no significant differences regarding PRA between the groups. There was a near-significant difference in cold ischemia time with longer times being observed for the ACR group (739±295 vs. 637±302; P=0.06; Mann-Whitney U test).

**Table 1.**
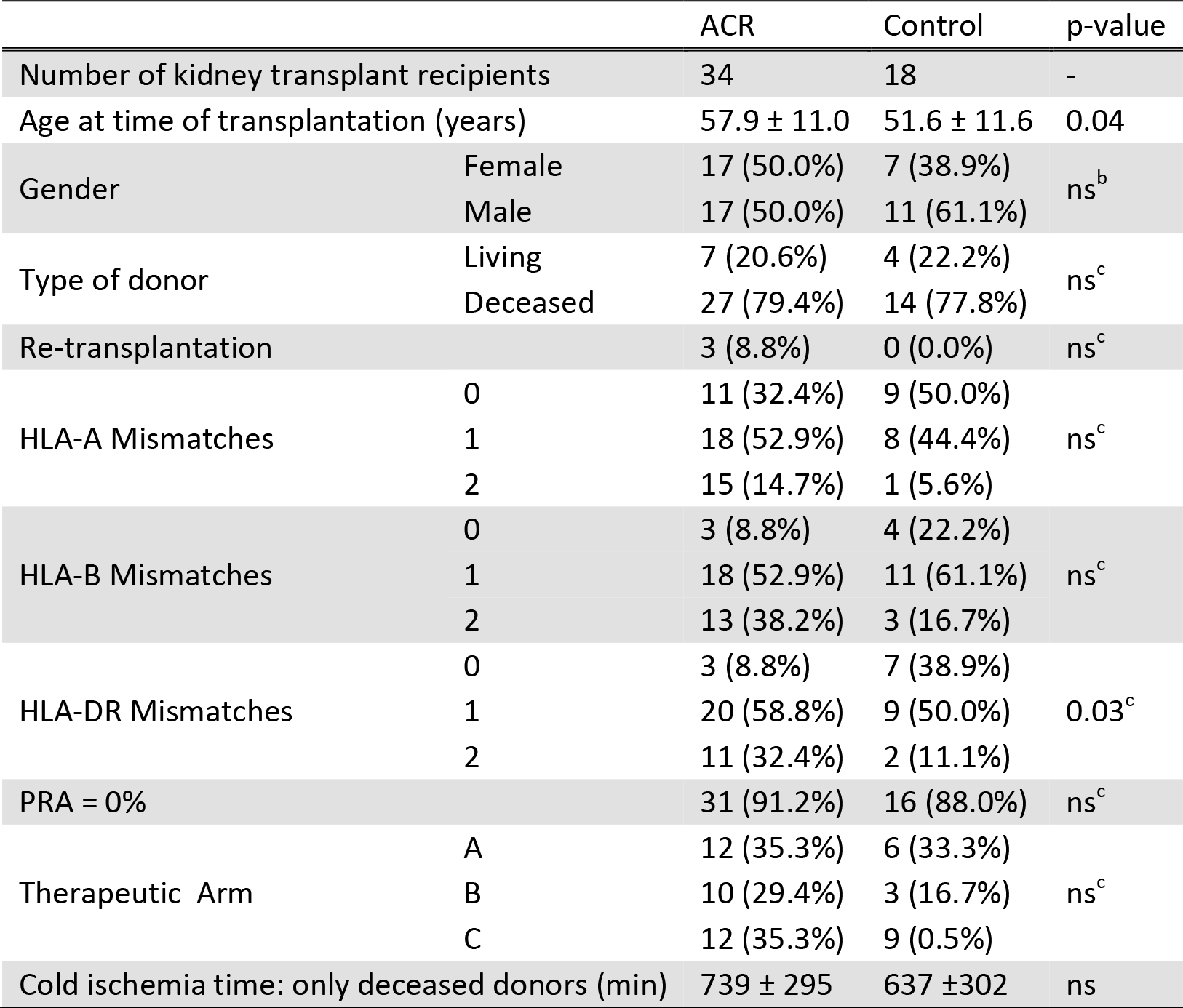
Characteristics and medication details for the subset of patients included in the HLA class 1 SAB data set^a^.

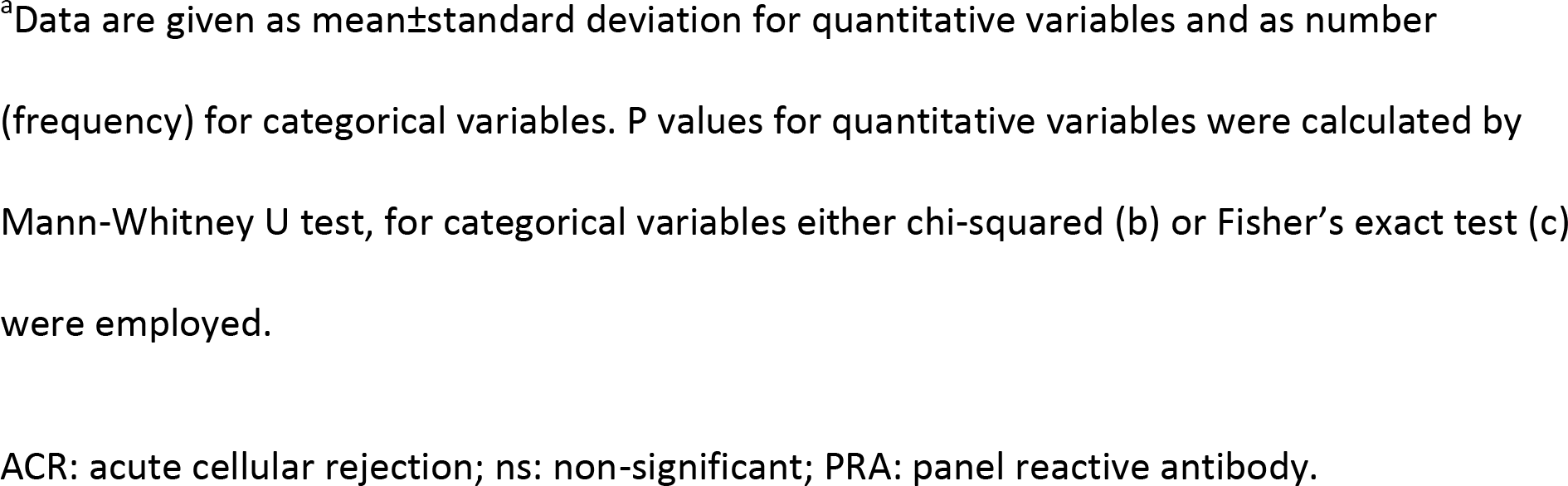

### 3.2. Conventional HLA SAB data analysis does not permit pre-transplant risk assessment of ACR

To assess whether HLA-1 SAB reactivity profiles provide a means for ACR risk assessment, we initially applied the conventional data analysis approach used in HLA-diagnostics to our data set. Central to this approach is the conversion of the quantitative SAB assay read-out data into qualitative binary data (1=presence of antibody-antigen reactivity, 0=absence of reactivity) based on a mean fluorescence intensity (MFI) threshold. We performed all analyses for a fixed threshold of 1000 MFI and an individually adjusted threshold in the range 253-1068 MFI (Supplemental Figure 2). In both cases, there were no statistically significant differences between the SAB ACR and the SAB control group in any of the individual reactivities (Supplemental Table 3 and 4). That is, there are no individual HLA-1 antibody reactivities that allow for risk assessment of ACR.

To assess whether there is a combination of reactivities that allows for risk assessment of ACR, we extended the conventional approach by applying a support vector machine-based multiparameter classification method to the binarized data (for details, see Material and Methods). The resulting multiparameter classifiers did not achieve significant classification performance (p>0.1, Table 2). Taken together, our results indicate that the conventional HLA SAB data analysis approach does not permit pre-transplant risk assessment of ACR.

**Table 2:** Multiparameter pre-transplant prediction of ACR^a^.

| Data set | Data analysis approach | MFI-treshold | BACC [%] | Sens. [%] | Spec. [%] | p-value |
| --- | --- | --- | --- | --- | --- | --- |
| HLA-1 SAB | Conventional, binary data input [0, 1] | 1000 | 62.1 | 35.3 | 88.9 | ns ($p>0.1$) |
| | Conventional binary data input [0, 1] | Individually adjusted [253-1068] | 70.9 | 52.9 | 88.9 | ns ($p>0.1$) |
|  | Novel, continuous data input | -- | 82.7 | 76.5 | 88.9 | 0.002 |
| MAB | Novel, continuous data input | -- | 63.9 | 55.6 | 72.2 | 0.040 |

<sup>a</sup>BACC: Balanced accuracy; MAB: mixed antigen bead screening; SAB: single antigen bead; Sens: Sensitivity; Spec: Specificity.

### 3.3. A novel approach built on multiparameter classification and quantitative data input allows for high accuracy pre-transplant prediction of ACR

In spite of the widespread use of HLA SAB assays, the interpretation of results obtained following the conventional data analysis approach remains controversial (47). A strict MFI threshold consistently identifying clinically relevant antibody-antigen reactivities is challenging to define (48). Since it is likely that the choice of MFI threshold compromises classification efforts, we applied a novel approach to the HLA-1 SAB data set that does not rely on MFI-thresholding. Key to this novel approach is the rank-normalization of the continuous SAB assay read-out data. Remarkably, a support-vector machine-based multiparameter classificator built on these data achieved highly significant prediction performance with a balanced accuracy of 82.7% (sensitivity=76.5%, specificity= 88.9%, p=0.002, Figure 1 A and Table 2). Receiver operating characteristic (ROC) analysis further emphasizes that the prediction performance was better than a random guess (area under the curve [AUC] =0.86) and illustrates the trade-off between the probability of correctly predicting ACR (true positive rate, sensitivity) and the probability of incorrectly predicting ACR (false positive rate, specificity) (Figure 1 B).

**Figure 1.**
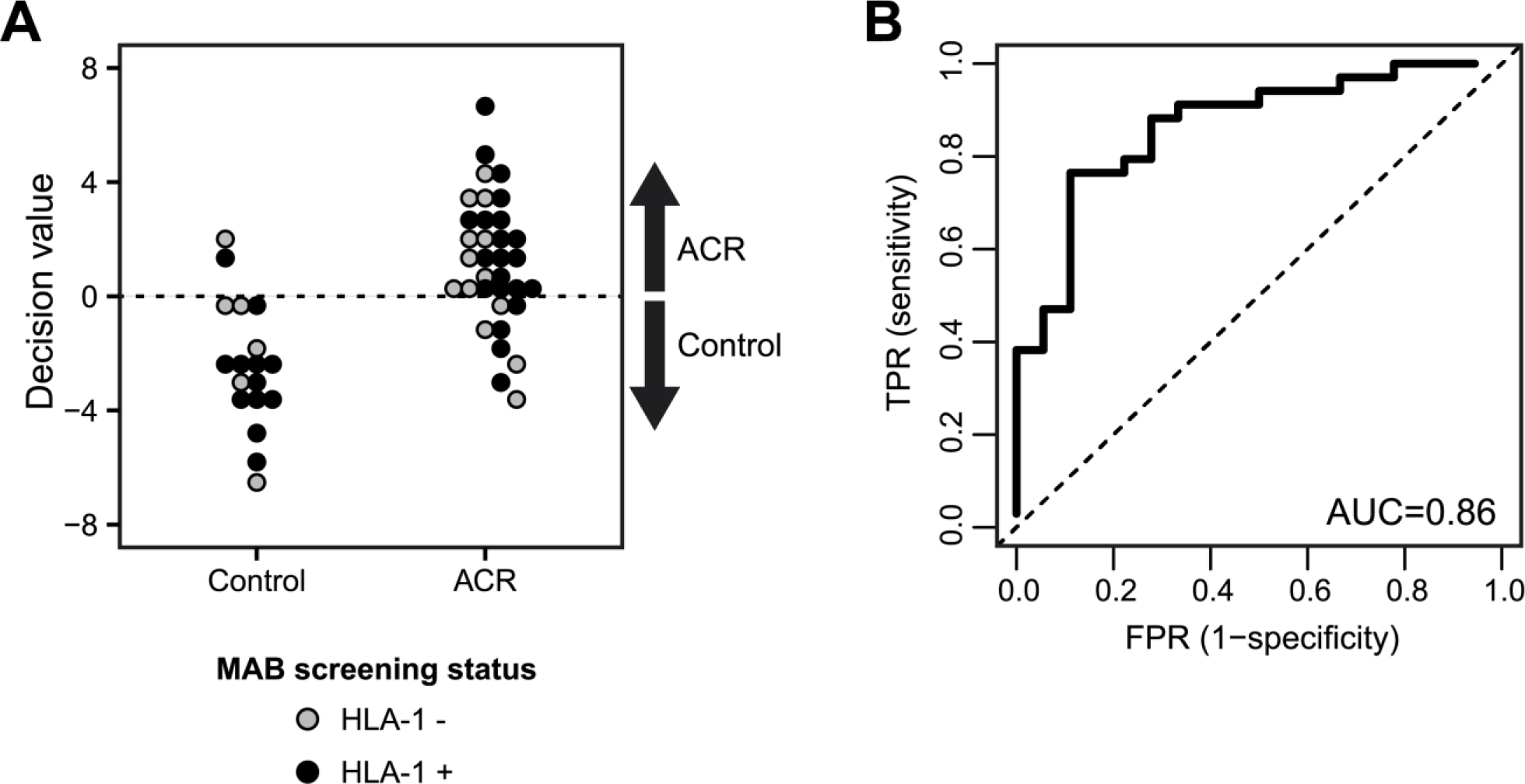
Predictive performance of the multiparameter ACR risk assessment classifier based on rank-normalized continuous pre-transplant HLA-1 antibody reactivity profiles. (A) Output of the classifiers decision function for each patient. The decision threshold is indicated by a dashed horizontal line. Patients with a decision value >0 are classified as ACR, patients with a decision value <0 are classified as control. Colors indicate whether patients tested positive (black) or negative (grey) for the presence of serum HLA-1 antibodies during MAB screening. (B) ROC curve of the multiparameter classifier. ACR: acute cellular rejection; ROC: receiver operating characteristic; SAB: single antigen bead screening; FPR: false positive rate; TPR: true positive rate.

Importantly, we found that prediction performance was independent of a patient’s MAB screening test result, as patients who tested positive or negative for HLA-1 antibodies are predicted equally well (Figure 1 A). Moreover, the performance of the continuous data classifier was not due to age or HLA-DR mismatch frequency as confounding factors; significant classification was not achieved when HLA class 1 SAB continuous data were grouped according to either of those factors (≤50 y vs. >50 y or no-mismatch vs. 1-2 mismatches). In addition, median-centered bead MFIs did not show any association with age (mean Pearson correlation coefficient r=0.019 ± 0.129). Taken together, our results show that continuous, rank-normalized HLA-1 SAB reactivity profiles provide a means of high-accuracy risk assessment of pre-transplant ACR.

### 3.4. Diagnostics based on HLA antibody detection assays may generally benefit from the novel approach

The fact that continuous HLA-1 SAB reactivity data outperformed MFI-thresholded binary data in terms of pre-transplant prediction of ACR (Table 2) led us to the conjecture that the conventional approach entails a loss of information that may compromise HLA-diagnostics classification efforts in general. To substantiate this claim, we performed additional analyses on the MAB screening data (63 ACR + 54 controls). Conventional MFI-threshold based data analysis revealed no statistical differences between the two study group as to the prevalence of HLA class 1 and/or HLA class 2 antibodies (Figure 2). A multiparameter classifier based on the continuous rank-normalized data, however, achieved statistically significant prediction of the patients who experience ACR (p=0.04, Table 2 and Supplemental Figure 3). Even though the accuracy of the classifier was low and not sufficient for routine risk assessment (balanced accuracy=63.9%, sensitivity=55.6%, specificity= 72.2%), the fact that it was significant emphasizes that the use of continuous non-thresholded antigen bead assay data favorably affects classification performance.

**Figure 2:**
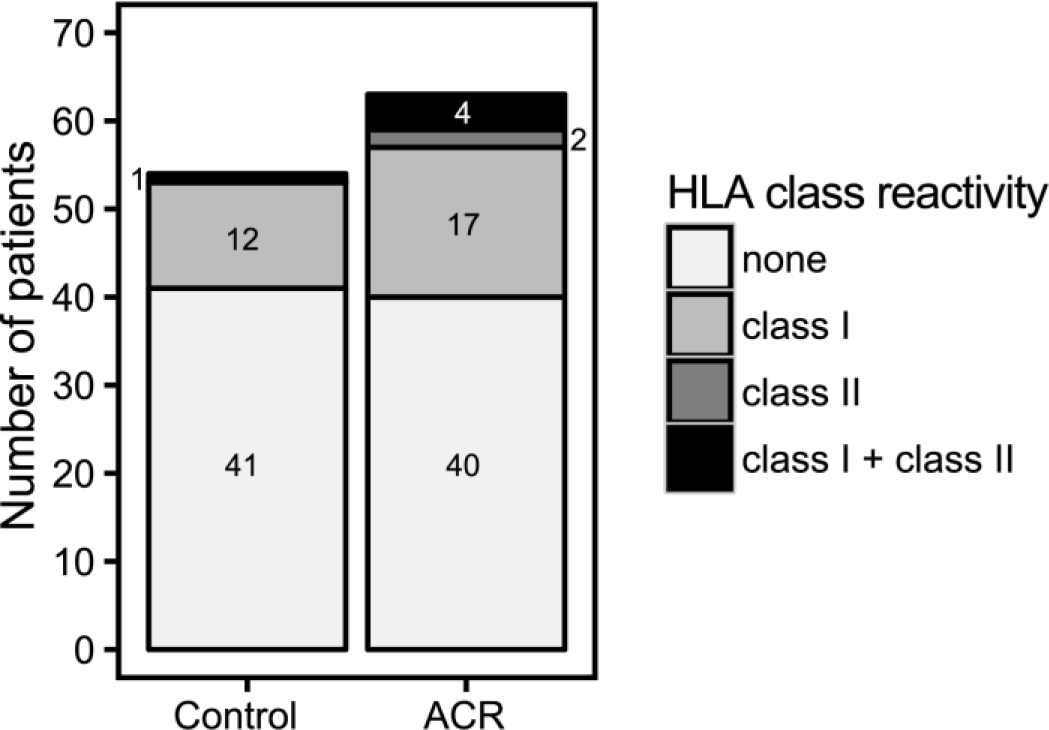
Conventional MFI-thresholded binary MAB screening data do not allow for pre-transplant risk assessment of ACR. Illustrated are the results of the MAB screening data of the cohort (117 graft recipients, 63 ACR + 54 controls; for demographics and clinical characteristics, see Supplemental Table 2).). Analyses were carried out on MFI-thresholded binary HLA MAB screening data (conventional approach); according to Fisher’s exact test, differences with respect to the prevalence of HLA antibodies are not significant (p>0.05). ACR: acute cellular rejection; MAB: mixed antigen bead screening.

## 4. Discussion

The current study shows that pre-transplant HLA class 1 SAB signatures predict the risk of acute cellular rejection (ACR) with high accuracy. Importantly, it demonstrates that HLA antibody signatures contain information on cell-mediated events to come.

In contrast to the vast majority of existing pre-transplant risk assessment models (22–27,31), our model does not rely on DSA reactivity data. The key advantages of this approach are that i) it facilitates risk assessment for non-sensitized patients lacking DSA and ii) it can be carried out independently of donor assignment.

HLA SAB data usually feed into prediction models in the form of binary data derived from MFI thresholding – the focus lying on strong binding interactions. Strikingly, our study emphasizes that such an approach entails a loss of information and ultimately results in loss of or suboptimal prediction performance. The fact that continuous HLA reactivity data outperform thresholded binary data (Table 2) indicates that weak binding interactions hold high-value information for risk assessment of ACR. This is further emphasized by our finding that our risk assessment tool performs equally well for patients who tested positive or negative for the presence of HLA-1 antibodies during MAB screening. There is indeed sufficient evidence in the literature to show that weak binding events are of great importance to biological systems, e.g. protein–peptide interactions (49), virus-cell interactions (50), cell adhesion, and cell–cell interactions (51–54). Our data suggest that HLA SAB based diagnostics will profit from inclusion of weak interactions by feeding prediction models with non-thresholded continuous data. A further advantage of prediction models based on non-thresholded MFI data is that they are not affected by the prevailing uncertainties regarding the right choice of the threshold MFI level, and yet missing internationally agreed standards (47,55).

But why is the pre-transplant signature of serum antibodies against HLA-1 SAB predictive for the risk of T cell mediated rejection? There is evidence in the literature for an association between anti-HLA serum antibodies and ACR. However, the nature of our classifier is as yet elusive, as it is commonly the case for “big data” multiparameter diagnostic approaches. Immunosignaturing effects have been described in the literature as the capacity of serum antibody binding profiles against protein or peptide libraries to discriminate between health and disease conditions. For instance, immunosignatures have been shown to aid fine diagnostics of brain tumors, the early diagnosis of Alzheimer’s disease as well as assessment of vaccine efficacy (36–39).

The HLA MAB and SAB data used in this study are part of a large multi-parameter database set up by the e:KID consortium that seeks to establish a systems medicine approach to personalized immunosuppressive treatment at an early stage after kidney transplantation (http://www.sys-med.de/en/consortia/ekid/). e:KID recorded a total of 478 parameters including, among others, gene expression, cytokine profile, epigenetics, metabolomics and viral load data as well as common clinical variables such as renal function or acute phase proteins. Evaluation of clinical parameters failed to identify any markers or combinations thereof which are predictive of ACR (4). Additionally, no other single parameter or multiparameter set, other than HLA class 1 SAB signatures, achieved high accuracy pre-transplant prediction performance. This emphasizes the vast potential of serum antibodies in diagnostics in general, and, in particular, for diseases where the antigens are unknown.

The comparison of predictive performances between our classifier and classifiers in the literature underlines its relevance to pre-kidney transplant risk assessment (Supplemental Table 5). Its accuracy of 82.7% is i) one of the highest among all donor-independent risk assessment models (19,25,30,34,57), ii) comparable to any AR models (19,20,30,34,35,57,58,21,22,24–29) and iii) comparable to any SAB data based models for ABMR (22,24–28). Furthermore, our classifier is based on SAB, an established diagnostics laboratory tool, thereby facilitating its further use for ACR risk assessment.

In conclusion, our study establishes a novel tool for pre-transplant risk assessment of acute cellular rejection. Once externally validated, patients classified as high risk by our model will benefit from its implementation through modified immunosuppression as well as closer monitoring leading to earlier detection of rejection onset and initiation of treatment. Consequently, the prognosis and survival rate of the graft will improve.

## Acknowledgements

This work was supported by the German Federal Ministry of Education and Research (BMBF) within the framework of the e:Med research and funding concept (01ZX1312). The authors thank Dr. Nils Lachmann, Center for Tumor Medicine, H&I Laboratory, Charité University Medicine Berlin, for helpful advice and discussions.

## Description of Supporting Information

Additional Supporting Information may be found online in the supporting information tab for this article.

**Supplemental Figure 1. Cumulative frequency of ACR through the first post-transplantation year.** Frequency of patients in the ACR group who experienced at least one ACR event at different time points post-transplantation. At week 2 and month 3, 34% and 79% of patients in the ACR group had experienced an ACR event.

ACR: acute cellular rejection

**Supplemental Figure 2. Distribution of MFI cutoffs for generation of binary HLA class 1 SAB data.** Illustrated are the MFI cutoff values of the 52 pre-transplant serum samples (control group=18; ACR group, Rej=34) of the binary HLA class 1 SAB data (1: presence of reactivity=above the MFI cutoff).

MFI: mean fluorescence intensity; SAB: single antigen bead screening.

**Supplemental Figure 3. Predictive performance of multiparameter ACR classification based on rank-normalized continuous pre-transplant MAB screening data. (**A) Output of the classifiers decision function for each patient. The decision threshold is indicated by a dashed horizontal line. Patients with a decision value >0 are classified as ACR, patients with a decision value <0 are classified as control. Colors indicate whether patients tested positive (black) or negative (grey) for the presence of serum HLA-1 antibodies during MAB screening. (B) ROC curve of the multiparameter classifier.

ACR: acute cellular rejection; ROC: receiver operating characteristic; SAB: single antigen bead screening; FPR: false positive rate; TPR: true positive rate.

**Supplemental Table 1.** Baseline characteristics and medication details for the Harmony patient cohort (N=157).^a^

ACR: acute cellular rejection; ns: non-significant; PRA: panel reactive antibody.

**Supplemental Table 2.** Baseline characteristics and medication details for the patients with HLA MAB data (N=117).^a^

ACR: acute cellular rejection; ns: non-significant; MAB: mixed HLA antigen bead; PRA: panel reactive antibody.

**Supplemental Table 3.** Single parameter prediction of ACR based on binarized HLA class 1 SAB data (fixed MFI threshold of 1000 MFI).

ACR: acute cellular rejection; ns: non-significant; SAB: single antigen bead screening; PRA: panel reactive antibody.

**Supplemental Table 4.** Single parameter prediction of ACR based on binarized HLA class 1 SAB data (individually adjusted MFI threshold).

ACR: acute cellular rejection; ns: non-significant; SAB: single antigen bead screening; PRA: panel reactive antibody.

**Supplemental Table 5.** Summary of existing AR prediction models. SAB: single antigen bead screening.

## Abbreviations

ABMR: acute antibody-mediated rejection
ACR: acute cellular rejection
AR: acute rejection
AUC: area under the curve
BACC: balanced accuracy
DSA: donor-specific antibodies
IQR: interquartile range
KTR: kidney transplantation
MAB: mixed HLA antigen bead
MFI: median fluorescence intensity
PRA: Panel Reactive Antibodies
P-SVM: Potential Support Vector Machine
pretransplantation: pre-Tx
posttransplantation: post-Tx
ROC: receiver operating characteristic
SAB: single HLA antigen bead

## References

1. Djamali A, Kaufman DB, Ellis TM, Zhong W, Matas A, Samaniego M. Diagnosis and management of antibody-mediated rejection: Current status and novel approaches. Am J Transplant. (2014);14(2):255–71.

2. Meier-Kriesche HU, Schold JD, Srinivas TR, Kaplan B. Lack of Improvement in Renal Allograft Survival Despite a Marked Decrease in Acute Rejection Rates over the Most Recent Era. Am J Transplant. (2004);4(3):378–83.

3. Meier-Kriesche HU, Schold JD, Kaplan B. Long-term renal allograft survival: Have we made significant progress or is it time to rethink our analytic and therapeutic strategies? Am J Transplant. (2004);4(8):1289–95.

4. Thomusch O, Wiesener M, Opgenoorth M, Pascher A, Woitas RP, Witzke O, et al. Rabbit-ATG or basiliximab induction for rapid steroid withdrawal after renal transplantation (Harmony): an open-label, multicentre, randomised controlled trial. Lancet. (2016);388:3006–16.

5. Ciancio G, Burke GW, Gaynor JJ, Carreno MR, Cirocco RE, Mathew JM, et al. A Randomized Trial of Three Renal Transplant Induction Antibodies: Early Comparison of Tacrolimus, Mycophenolate Mofetil, and Steroid Dosing, and Newer Immune-Monitoring1. Transplantation. (2005);80(4):457–65.

6. Ekberg H, Tedesco-Silva H, Demirbas A, Vítko Š, Nashan B, Gürkan A, et al. Reduced Exposure to Calcineurin Inhibitors in Renal Transplantation. N Engl J Med. (2007);357(25):2562–75.

7. El-Zoghby ZM, Stegall MD, Lager DJ, Kremers WK, Amer H, Gloor JM, et al. Identifying specific causes of kidney allograft loss. Am J Transplant. (2009);9(3):527–35.

8. Roberts DM, Jiang SH, Chadban SJ. The Treatment of Acute Antibody-Mediated Rejection in Kidney Transplant Recipients—A Systematic Review. Transplant J. (2012);94(8):775–83.

9. Magil AB. Monocytes/macrophages in renal allograft rejection. Transplant Rev. (2009);23(4):199–208.

10. Becker LE, Morath C, Suesal C. Immune mechanisms of acute and chronic rejection. Clin Biochem. (2016);49(4-5):320–3.

11. Suthanthiran M, Schwartz JE, Ding R, Abecassis, Michael Dadhania D, Samstein B, Knechtle SJ, et al. Urinary-Cell mRNA Profile and Acute Cellular Rejection in Kidney Allografts. N Engl J Med. (2013);369:20–31.

12. Afaneh C, Muthukumar T, Lubetzky M, Ding R, Snopkowski C, Sharma VK, et al. Urinary cell levels of mRNA for OX40, OX40L, PD-1, PD-L1, or PD-L2 and acute rejection of human renal allografts. Transplantation. (2010);90(12):1381–7.

13. Hricik DE, Nickerson P, Formica RN, Poggio ED, Rush D, Newell KA, et al. Multicenter validation of urinary CXCL9 as a risk-stratifying biomarker for kidney transplant injury. Am J Transplant. (2013);13(10):2634–44.

14. Roedder S, Sigdel T, Salomonis N, Hsieh S, Dai H, Bestard O, et al. The kSORT Assay to Detect Renal Transplant Patients at High Risk for Acute Rejection: Results of the Multicenter AART Study. PLoS Med. (2014);11(11).

15. Wang JN, Zhou Y, Zhu TY, Wang X, Guo YL. Prediction of acute cellular renal allograft rejection by urinary metabolomics using MALDI-FTMS. J Proteome Res. (2008);7(8):3597–601.

16. Reeve J, Einecke G, Mengel M, Sis B, Kayser N, Kaplan B, et al. Diagnosing rejection in renal transplants: A comparison of molecular- and histopathology-based approaches. Am J Transplant. (2009);9(8):1802–10.

17. Desvaux D, Schwarzinger M, Pastural M, Baron C, Abtahi M, Berrehar F, et al. Molecular diagnosis of renal-allograft rejection: correlation with histopathologic evaluation and antirejection-therapy resistance. Transplantation. (2004);78(5):647–53.

18. Ting YT, Coates PT, Marti H-P, Dunn AC, Parker RM, Pickering JW, et al. Urinary soluble HLA-DR is a potential biomarker for acute renal transplant rejection. Transplantation. (2010);89(9):1071–8.

19. Poggio ED, Augustine JJ, Clemente M, Danzig JM, Volokh N, Zand MS, et al. Pretransplant Cellular Alloimmunity as Assessed by a Panel of Reactive T Cells Assay Correlates With Acute Renal Graft Rejection. Transplantation. (2007);83(7):847–52.

20. Simon T, Opelz G, Wiesel M, Pelzl S, Ott RC, Süsal C. Serial Peripheral Blood Interleukin-18 and Perforin Gene Expression Measurements for Prediction of Acute Kidney Graft Rejection. Am J Transplant. (2003);3(1121-1127):1589–95.

21. Dong W, Shunliang Y, Weizhen W, Qinghua W, Zhangxin Z, Jianming T, et al. Prediction of acute renal allograft rejection in early post-transplantation period by soluble CD30. Transpl Immunol. (2006);16(1):41–5.

22. Malheiro J, Tafulo S, Dias L, Martins LS, Fonseca I, Beirão I, et al. Analysis of preformed donor-specific anti-HLA antibodies characteristics for prediction of antibody-mediated rejection in kidney transplantation. Transpl Immunol. (2015);32(2):66–71.

23. Vlad G, Ho EK, Vasilescu ER, Colovai AI, Stokes MB, Markowitz GS, et al. Relevance of different antibody detection methods for the prediction of antibody-mediated rejection and deceased-donor kidney allograft survival. Hum Immunol. (2009);70(8):589–94.

24. Riethmüller S, Ferrari-Lacraz S, Müller MK, Raptis D a., Hadaya K, Rüsi B, et al. Donor-Specific Antibody Levels and Three Generations of Crossmatches to Predict Antibody-Mediated Rejection in Kidney Transplantation. Transplant J. (2010);90(2):160–7.

25. Lefaucheur C, Loupy A, Hill GS, Andrade J, Nochy D, Antoine C, et al. Preexisting donor-specific HLA antibodies predict outcome in kidney transplantation. J Am Soc Nephrol. (2010);21(8):1398–406.

26. Song EY, Lee YJ, Hyun J, Kim YS, Ahn C, Ha J, et al. Clinical relevance of pretransplant HLA Class II Donor-specific antibodies in renal transplantation patients with negative T-cell cytotoxicity crossmatches. Ann Lab Med. (2012);32(2):139–44.

27. Salvadé I, Aubert V, Venetz JP, Golshayan D, Saouli AC, Matter M, et al. Clinically-relevant threshold of preformed donor-specific anti-HLA antibodies in kidney transplantation. Hum Immunol. (2016);77(6):483–9.

28. Shaikhina T, Lowe D, Daga S, Briggs D, Higgins R, Khovanova N. Decision tree and random forest models for outcome prediction in antibody incompatible kidney transplantation. Biomed Signal Process Control. (2017);1–7.

29. Hauser IA. Prediction of Acute Renal Allograft Rejection by Urinary Monokine Induced by IFN-(MIG). J Am Soc Nephrol. (2005);16(6):1849–58.

30. Mancebo E, Castro MJ, Allende LM, Talayero P, Brunet M, Millán O, et al. High proportion of CD95+ and CD38+ in cultured CD8+ T cells predicts acute rejection and infection, respectively, in kidney recipients. Transpl Immunol. (2016);34:33–41.

31. Ho EK, Vasilescu ER, Colovai AI, Stokes MB, Hallar M, Markowitz GS, et al. Sensitivity, specificity and clinical relevance of different cross-matching assays in deceased-donor renal transplantation. Transpl Immunol. (2008);20(1-2):61–7.

32. Kannabhiran D, Lee J, Schwartz JE, Friedlander R, Aull M, Muthukumar T, et al. Characteristics of Circulating Donor Human Leukocyte Antigen-specific Immunoglobulin G Antibodies Predictive of Acute Antibody-mediated Rejection and Kidney Allograft Failure. Transplantation. (2015);99(6):1156–64.

33. Poggio ED, Clemente M, Hricik DE, Heeger PS. Panel of Reactive T Cells as a Measurement of Primed Cellular Alloimmunity in Kidney Transplant Candidates. J Am Soc Nephrol. (2006);17:564–72.

34. Vondran FWR, Timrott K, Kollrich S, Steinhoff AK, Kaltenborn A, Schrem H, et al. Pre-transplant immune state defined by serum markers and alloreactivity predicts acute rejection after living donor kidney transplantation. Clin Transplant. (2014);28(9):968–79.

35. Nafar M, Farrokhi F, Vaezi M, Entezari AE, Pour-Reza-Gholi F, Firoozan A, et al. Pre-transplant and post-transplant soluble CD30 for prediction and diagnosis of acute kidney allograft rejection. Int Urol Nephrol. (2009);41(3):687–93.

36. Hughes AK, Cichacz Z, Scheck A, Coons SW, Johnston SA, Stafford P. Immunosignaturing Can Detect Products from Molecular Markers in Brain Cancer. PLoS One. 2012 Jul;7(7).

37. Restrepo L, Stafford P, Johnston SA. Feasibility of an early Alzheimer’s disease immunosignature diagnostic test. J Neuroimmunol. 2013 Jan;254(1-2):154–60.

38. Stafford P, Cichacz Z, Woodbury NW, Johnston SA. Immunosignature system for diagnosis of cancer. Proc Natl Acad Sci U S A. 2014 Jul;111(30):E3072–3080.

39. Legutki JB, Johnston SA. Immunosignatures can predict vaccine efficacy. Proc Natl Acad Sci U S A. 2013 Nov;110(46):18614–9.

40. Cascalho MI, Chen BJ, Kain M, Platt JL. The paradoxical functions of B cells and antibodies in transplantation. J Immunol. (2013);190(3):875–9.

41. Randhawa P. T-cell-mediated rejection of the kidney in the era of donor-specific antibodies: diagnostic challenges and clinical significance. Curr Opin Organ Transplant. (2015);20(3):325–32.

42. Solez K, Colvin RB, Racusen LC, Sis B, Halloran PF, Birk PE, et al. Banff’05 meeting report: Differential diagnosis of chronic allograft injury and elimination of chronic allograft nephropathy (‘CAN’). Am J Transplant. (2007);7(3):518–26.

43. Blazquez-Navarro A, Schachtner T, Stervbo U, Sefrin A, Stein M, Westhoff TH, et al. Differential T cell response against BK virus regulatory and structural antigens: A viral dynamics modelling approach. PLOS Comput Biol. (2018);14(5):1–20.

44. Blazquez-Navarro A, Dang-Heine C, Wittenbrink N, Bauer C, Wolk K, Sabat R, et al. BKV, CMV, and EBV Interactions and their Effect on Graft Function One Year Post-Renal Transplantation: Results from a Large Multi-Centre Study. EBioMedicine. (2018);

45. Molinaro AM, Simon R, Pfeiffer RM. Prediction error estimation: A comparison of resampling methods. Bioinformatics. (2005);21(15):3301–7.

46. Mason SJ, Graham NE. Areas beneath the relative operating characteristics (ROC) and relative operating levels (ROL) curves: Statistical significance and interpretation. Q J R Meteorol Soc. (2002);128(584):2145–66.

47. Sullivan HC, Liwski RS, Bray RA, Gebel HM. The Road to HLA antibody evaluation: Do not rely on MFI. Am J Transplant. (2017);XX:1–7.

48. Konvalinka A, Tinckam K. Utility of HLA Antibody Testing in Kidney Transplantation. J Am Soc Nephrol. (2015);26(7):1489–502.

49. Fairchild PJ, Wraith DC. Lowering the tone: Mechanisms of immunodominance among epitopes with low affinity for MHC. Immunol Today. (1996);17(2):80–5.

50. Haywood AM. Virus receptors: binding, adhesion strengthening, and changes in viral structure. J Virol. (1994);68(1):1–5.

51. Hakomori S. Structure and Function of Sphingoglycolipids in Transmembrane Signalling and Cell-Cell Interactions. Biochem Soc Trans. (1993);21(3):583–95.

52. van der Merwe PA, Brown MH, Davis SJ, Barclay AN. Affinity and kinetic analysis of the interaction of the cell adhesion molecules rat CD2 and CD48. EMBO J. (1993);12(13):4945–54.

53. Reilly PL, Woska JR, Jeanfavre DD, McNally E, Rothlein R, Bormann BJ. The native structure of intercellular adhesion molecule-1 (ICAM-1) is a dimer. Correlation with binding to LFA-1. J Immunol. (1995);155(2):529–32.

54. Hage DS. Weak Affinity Chromatography. In: Affinity Chromatography Methods and Protocols. 2000. p. 7–23.

55. Szatmary P, Jones J, Hammad A, Middleton D. Impact of sensitivity of human leucocyte antigen antibody detection by Luminex technology on graft loss at 1 year. Clin Kidney J. (2013);6(3):283–6.

56. Lobo LJ, Aris RM, Schmitz J, Neuringer IP. Donor-specific antibodies are associated with antibody-mediated rejection, acute cellular rejection, bronchiolitis obliterans syndrome, and cystic fibrosis after lung transplantation. J Hear Lung Transplant. 2013 Jan;32(1):70–7.

57. Pike R, Thomas N, Workman S, Ambrose L, Guzman D, Sivakumaran S, et al. PD1-expressing T cell subsets modify the rejection risk in renal transplant patients. Front Immunol. (2016);7(APR).

58. Zhang Q, Liu YF, Su ZX, Shi LP, Chen YH. Serum fractalkine and interferon-gamma inducible protein-10 concentrations are early detection markers for acute renal allograft rejection. Transplant Proc. (2014);46(5):1420–5.

